# High-altitude mountaineering induces adaptive gut microbiome shifts associated with dietary intake and performance markers

**DOI:** 10.1101/2025.06.06.658263

**Authors:** Ewa Karpęcka-Gałka, Kinga Zielińska, Barbara Frączek, Paweł Łabaj, Tomasz Kościółek, Kinga Humińska-Lisowska

## Abstract

This study examined how high-altitude exposure and expedition-specific dietary changes influence gut microbiome composition, functional pathways, and their relationships with performance and health markers in alpinists. Seventeen male mountaineers (age 30.29±5.8 years) participating in multi-week expeditions (>3,000 MASL) were assessed before and after their climbs. Assessments included dietary intake analysis, blood and urine biomarkers, aerobic and anaerobic performance tests, and metagenomic sequencing of the gut microbiome. Bioinformatic and statistical analyses evaluated changes in microbiome composition and function and their correlations with physiological and dietary parameters. High-altitude exposure was associated with significant shifts in gut microbial composition and functional capacity. While the total number of bacterial species and functions remained stable, the glucose degradation pathway increased post-expedition. Participants with greater microbiome shifts showed improved performance and had richer baseline microbiomes. Pre-expedition, certain microbial functions were associated with vitamin B_6_ and C intake, while post-expedition correlations involved specific macronutrients and micronutrients. Additionally, some microbiome changes correlated with blood markers, indicating links to nutrient metabolism and electrolyte balance. The gut microbiome of alpinists adapts to extreme environmental stress and dietary changes, influencing metabolic, immune, and performance-related processes. Optimizing dietary strategies to support a beneficial microbiome profile may enhance resilience and performance in challenging high-altitude environments.

## 1. Introduction

During climbing at high altitudes (>2,500 MASL), alpinists must deal with reduced atmospheric pressure, leading to the development of hypobaric hypoxia, intense solar radiation, variable weather conditions, strong winds and low temperatures with large fluctuations throughout the day, as well as the risk of avalanches and falling rocks^1,2^. During high-altitude expeditions, dietary changes occur due to poor access to fresh food, lack of appetite, food poisoning, harsh environmental conditions, and physiological changes^3,4^. Reactive oxygen species (ROS) production is accelerated with mountainous elevation, which may play a role in developing serious health crises. Exposure to increasing altitude leads to a reduction in ambient O_2_ availability in cells, contributing to hypoxic oxidative stress and disturbed redox homeostasis, potentially impairing exercise performance and recovery^5,6^. As a result of increased ROS production, alpinists may experience cognitive decline, development of neurodegenerative processes, pathological changes in brain structures, including vasogenic edema^7^, as well as damage to the intestinal barrier. This intestinal barrier disruption may lead to bacterial translocation and local or systemic inflammatory reactions, further impacting both general health and physical performance^8^. A damaged intestinal barrier may impair absorption of nutrients, which could explain some of the weight loss observed during prolonged exposures to high altitude^9,10^.

In addition to these physiological stresses, blood parameters and body composition can be altered by high-altitude conditions and changes in dietary intake. This may affect red blood cell counts, iron status, and markers of metabolic and immune function^11–15^. Similarly, adjustments in macronutrient and micronutrient intake before and during the expedition can affect an athlete’s energy availability, leading to shifts in body mass and composition^16^, and potentially affecting VO_2max,_ anaerobic power, and other key performance indicators. Such high-altitude exposures thus serve as a natural experimental setting to assess how nutrition, physiology, and microbial ecology interact to shape athletic performance, health status, and adaptation to extreme environments.

In recent years, the gut microbiome has emerged as a critical factor influencing human health, immune function, and athletic performance. Alterations in gut microbial composition and metabolic capacity have been linked to changes in nutrient absorption, energy metabolism, inflammation, and recovery, all of which are central to optimal athletic performance and well-being^17–19^. In addition, the gut microbiota can respond dynamically to dietary changes, and high-altitude expeditions often require consumption of processed, freeze-dried, or supplemental foods due to limited access to fresh produce^20^. Understanding how these dietary and environmental stresses shape gut microbial communities is essential for informed nutritional strategies that support health, performance, and safe acclimatization to challenging conditions.

The present study aimed to analyze the structure and functional characteristics of the microbiome of climbers before and after a high-mountain expedition. By integrating detailed dietary records, blood and urine analyses, and measures of aerobic and anaerobic exercise capacity, we sought to determine how high-altitude exposure and associated dietary changes affect the gut microbiome and, in turn, how these microbial shifts correlate with health markers and performance outcomes. Such insights could provide evidence-based guidance for optimizing dietary composition, supplementation, and overall expedition planning. Ultimately, identifying specific microbial signatures and metabolic pathways that are modifiable through diet and supplementation may help practitioners develop tailored nutritional interventions to improve both health and athletic performance of climbers operating under extreme environmental conditions.

## 2. Materials and methods

### 2.1. Study Participants

The study group consisted of 17 male mountaineers from Poland between the ages of 23 and 40, participating in mountain expeditions at least once a year, during which they stayed at altitudes above 3,000 meters for at least 3 weeks. Individuals qualified for the study were members of high-mountain clubs affiliated with the Polish Mountaineering Association. The group’s anthropometric and BMI (body mass index) data are shown in Table 1.

**Table 1.** Anthropometric data and BMI of climbers participating in the research (n = 17)

|  | Men (n=17) |  |
| --- | --- | --- |
|  | value | SD |
| Age [years] | 30.29 | 5.8 |
| Body height [cm] | 180.47 | 8.36 |
| Body weight [kg] | 74.96 | 5.03 |
| BMI [kg/m<sup>2</sup>] | 22.83 | 2.1 |

Abbreviations: BMI - body mass index.

The climbers participating in the study were characterized by extensive climbing experience (10 ± 5 years) in both sport climbing and mountaineering. The average climbing level of participants, as measured by the IRCRA scale, was 20.69 ± 2.8, which according to the proposed criteria, places them in the advanced (level 3) category^21^. The climbers spent an average of 8 ± 3 hours per week training. Climbers qualified for the study stayed at an altitude of 3,000-8,167 meters above sea level at the time of their expedition. In total, the high-altitude expeditions lasted an average of 34 ± 6 days, of which climbers spent an average of 22 ± 4 days actively climbing in the mountains and 11 ± 8 days resting in base camps or mountain towns. Because some of these sites were located above 3,000 m, participants spent a total of 26 ± 6 days at altitudes above 3,000 m.

The mountaineers recruited for the study traveled in small groups or individually to pursue their mountain goals. Specifically, the objective of the group of climbers (n=6) exploring the White Cordillera massif in Peru was to climb the 800-meter Cruz del Sur route on the La Esfinge rock monolith (5,325 m). A new route was also established on Ocschapalca (5,888 m) and Nevado Churup (5,495 m), and ascents were made on Artesonraju (6,025 m) and Alpamayo (5,947 m). Climbers (n=3) targeting peaks from the Shuijerab mountain group in North Karakorum, Pakistan, made a route on the west face of the virgin peak of Trident Peak (6,150 m), and climbed the virgin peak of Sakwa Sar (6,050 m). Climbers (n=3) exploring the Gangotria Valley in India’s Garhwal Himalaya and mountaineers (n=2) climbing in the Himalayas in Western Nepal and Northern Nepal attempted virgin peaks, but failed due to weather conditions (the highest point of the expeditions - 5,000 meters). There were also two independently active climbers in the Himalayas - one of them made ascents of Annapurna (8,091 m) and Dhaulagiri (8,167 m) in Central Nepal, while the other climbed Ama Dablam (6,812 m). In the western Pamir-Alay in Kyrgyzstan’s Lailak Valley, an ascent was made by the Troschenko route on the north face of Ak-su (5,217 m).

Before taking part in the project, the climbers had a consultation with a sports medicine doctor. The climbers’ fitness for mountaineering and participation in the project was assessed based on previous examinations (ECG, blood and urine tests). The presence of chronic diseases and age over 45 were disqualifying factors for participation in the study. The Bioethics Committee of the Cracow Regional Medical Chamber approved the research project (68/KBL/OIL/2022; date: 11/04/2022), which was conducted in accordance with the Declaration of Helsinki. After learning about the risks and benefits of taking part in the project, all climbers gave written informed consent to participate in this study.

### 2.2. Study Design

#### 2.2.1. Nutritional Analysis of Diet

The supply of selected macronutrients before and during the mountaineering expedition was determined by analyzing the whole-day rations obtained using the 3-day food diary method. The expedition participants were asked to note down all the foods, meals and supplements eaten and fluids drunk during the 3 days before the expedition (2 days of sports activity, 1 day of rest) and 3 days during the expedition (2 days of sports activity, 1 day of rest). In the mountain conditions, these were days of acclimatization and ascending towards the summit, and the activities noted during this time included mainly hiking with elements of climbing. Climbers were given detailed instructions on how to keep food diaries before leaving for the expedition in order to minimize recording errors. The diaries contained commercially available products, so the necessary data for analysis was read from labels and nutritional tables. In the case of food products such as bars, energy gels and freeze-dried products, climbers were asked to provide the product name, manufacturer and product weight.

The data from the diaries were meticulously entered into the Aliant Dietetic Calculator program (Anmarsoft, Gdańsk, Poland; version: 85; database: 6.2) for quantitative analysis of the climbers’ diets before and during the expedition. The program used the “Tables of food composition and nutritional value”^22^ as a database, taking into account food recipes and losses due to product processing.

#### 2.2.2. Anthropometric Measurements

Body height of participants was measured before the expedition using a Seca 217 anthropometer (Seca GmbH & Co. KG, Hamburg, Germany) with an accuracy of 1 mm. The climbers’ body weight was determined using InBody 120 body composition analyzer (Inbody Bldg., Seoul, Korea) in the morning after a standardized meal between 7:45 and 8:30 am.

#### 2.2.3. Health status analysis

For health status analysis, blood samples were taken from veins in the elbow pit to determine hematological and biochemical blood indices, i.e. peripheral blood count (hematocrit, hemoglobin concentration, erythrocyte volume (MCV), erythrocyte hemoglobin content (MCH), erythrocyte hemoglobin concentration (MCHC), erythrocyte anisocytosis index (RDW), erythrocyte count, platelets, platelet volume (MPV), white blood cell count - leukocytes, leukocyte differentiation: lymphocytes, monocytes, basophils, eosinophils, neutrophils), erythrocyte sedimentation rate (ESR), glucose, total cholesterol, high-density lipoprotein cholesterol (HDL), low-density lipoprotein cholesterol (LDL), triglycerides (TG), creatinine (Cr), urea (U), uric acid (UA), total protein, sodium, potassium, chloride, calcium, magnesium, phosphorus, iron, ferritin, vitamin D, vitamin B12, liver enzymes: ALT (alanine aminotransferase), AST (aspartate aminotransferase), acid phosphatase, GGTP (gamma- glutamyltranspeptydase), albumin and total bilirubin. The determinations were carried out in the morning and on an empty stomach, up to two weeks before the expedition and within two days (1.41 ± 0.69 days) after the expedition.

In addition to the blood determinations, a general urinalysis (specific gravity, pH, leukocytes, nitrites, protein, glucose, ketones, urobilinogen, bilirubin, blood (erythrocytes/hemoglobin), color, clarity) was performed, as well as a fecal examination (general fecal examination, test for parasites, lamblia, *Helicobacter pylori*, levels of zonulin and antitrypsin). Study participants were given a uniform procedure for collecting urine and stool samples on their own. After a plum-sized stool sample was gathered, it was placed in a tube and stored in a refrigerator (temp. 2-4° C) and then delivered to the laboratory within 24h after the collection. The samples of urine and feces were submitted for testing to the laboratory at the same time as blood was collected.

Measurements were made using flow cytometry (blood count), kinetic-photometric method (ESR), indirect potentiometry (sodium, potassium, chloride), spectrophotometry (total protein, calcium, phosphorus, magnesium, total cholesterol, HDL, triglycerides, U, UA, iron, AST, ALT, creatinine, total bilirubin, GGTP, glucose), direct chemiluminescence (vitamin B12, vitamin D, ferritin), strip tests (general urinalysis), immunochromatographic method with Giardia test from Hydrex Diagnostics (*Giardia lamblia* in feces), and by H. pylori Ag by nal von minden (*Helicobacter pylori* in feces) and microscopically (general fecal examination and evaluation of food debris, testing for parasites, lamblia). All laboratory analyses described here were performed using the following equipment: Alinity HQ (Abbott, Singapore), Roller 20 (Alifax, Italy), Alinity C (Abott, Singapore), Alinity I (Abott, Singapore) and Atellica 1500 (Siemens, Germany) and were commissioned as an external service, performed by Alab Laboratories (Warsaw, Poland).

#### 2.2.4. Aerobic and anaerobic performance testing

Aerobic capacity was assessed during a test with gradually increasing load performed until exhaustion, while anaerobic capacity was assessed during the Wingate test for the lower and upper limbs. Exercise tests were performed twice. The first test was performed about two weeks before the expedition, and the second test was performed up to four days (3.88 ± 1.49 days) after returning from the expedition at an altitude of about 383 meters above sea level.

##### 2.2.4.1. Graded Treadmill Test

The test was performed on a mechanical treadmill h/p/Cosmos Saturn (h/p/Cosmos Sports & Medical GmbH, Germany) according to the following scheme: 2-minute recording of baseline cardiorespiratory indices in a standing position, a 4-minute run at a constant speed of 8 km/h with a treadmill angle of 1°, followed by a 1.1 km/h increase in running speed every 2 minutes until a running speed of 16.8 km/h was reached. In the remaining part of the test, the running speed did not change and an increase in exercise intensity was obtained by raising the angle of the treadmill by 1° every 2 minutes. The test continued until exhaustion, subjective to the test participant. Respiratory indices were monitored continuously using a Cortex Metalyzer 3B ergospirometer (CORTEX Biophysik GmbH, Germany), heart rate recorded using a Polar S-410 cardiac monitor (Polar-Electro, Finland).

##### 2.2.4.2. Wingate anaerobic test

This test was performed in a sitting position without getting up from the saddle on a Monark 894e cycloergometer for the lower limbs (Monark Sports & Medical, Sweden) and Monark 834e for the upper limbs (Monark Sports & Medical, Sweden) in a modified 20- second version. The participants’ task was to perform crank turns at a maximally high rhythm throughout the test. Both tests were preceded by warm-ups, with the lower limb test lasting 5 minutes with a load of 100 watts and the upper limb test lasting 4 minutes with a load of 60 watts. The pedaling rhythm was 60 rpm. During the warm-up for the lower limb test, brief 5–second accelerations of the pedaling rhythm to about 80% of subjective maximum capacity occurred in 2^nd^, 4^th^ and 5^th^ minute and in the warm-up for the upper limb test in 2^nd^ and 4^th^ minute, respectively. After the warm-up, the participants performed stretching exercises for 2 to 3 minutes, after which the test proper was started. The test load was set at 7.5% and 4.5% of body weight in the lower and upper limb tests, respectively.

##### 2.2.4.3. Assessment of lactate concentration

Before the start (5 min before exercise) and at the end of all tests (3 and 20 min after exercise), blood was drawn from the fingertip capillaries into tubes containing a glycolysis inhibitor to determine plasma lactate concentration. Lactate concentration was determined using LC2389 reagent (Randox Laboratories Ltd., UK) by an enzymatic colorimetric method, which allows the determination of biochemical markers by using a specific enzymatic reaction that yields a colored product. The concentration of lactate was determined by the color change of the solution, assessed by reading the absorbance at 550 nm using a Thermo Scientific Nicolet Evolution 201 PC Control dual-beam UV/VIS spectrophotometer (Thermo Fisher Scientific, USA). The absorbance results were recorded electronically, and were then used to calculate the concentration of lactate in each sample.

#### 2.2.5. Microbiome analysis

##### 2.2.5.1. Sample collection

Stool was collected and immediately placed in stool storage containers, and these were placed in a Styrofoam box with a cooling insert. The box with the stool sample was kept in the refrigerator (temp. 2-4° C) and then transferred within 24h of collection to the project contractor. The stool was then placed in the freezer (temp. -80° C). The fecal samples were collected from participants in the same rounds as described in the section describing the health analysis (before and after the expedition).

##### 2.2.5.2. DNA isolation, quantitation and quantification

Approximately 200 mg of each frozen stool sample was processed for DNA isolation using the QIAamp PowerFecal Pro DNA Kit (Qiagen, Germany) following a modified protocol designed to enhance microbial cell disruption. This procedure combined physical, mechanical (bead-beating with a Bead Ruptor Elite; Omni International, USA), and chemical lysis steps. Homogenization involved short, high-speed cycles with cooling intervals. Key modifications included the following steps: after adding Solution CD1, the sample was incubated at 55°C for 10 minutes with 500 rpm shaking and vortexing every 2 minutes. Further, bead beating was performed for 15 seconds at 6 m/s, repeated 4 times with a 2- minute break after each cycle, followed by a short spin. Next, 40 µl of proteinase K was added, and the mixture was incubated at 25°C for 10 minutes. A final, 5th round of bead beating was performed for 15 seconds at 6m/s. The extracted DNA was assessed for concentration and purity using a NanoDrop spectrophotometer (Thermo Scientific, USA), and the integrity of the DNA was verified on 1% agarose gels. Only the DNA samples with sufficient quality and yield were selected for further processing. The DNA concentration was then re-evaluated and standardized using a Qubit 3.0 Fluorometer (Thermo Fisher Scientific, USA) and the Qubit DNA BR assay, ensuring accurate normalization prior to library preparation.

##### 2.2.5.3. Library preparation and sequencing

Metagenomic libraries were prepared from 100 ng of input DNA per sample, using the KAPA HyperPlus Library Preparation Kit (Roche, Basel, Switzerland) as described in our previous papers^23,24^. The fragmentation step was optimized to produce fragments with an average size of 300-500 bp. Indexed adapter ligation was performed using KAPA Universal dual-indexed adapters at a controlled concentration to maintain efficient ligation and minimize adapter dimer formation. Libraries were then purified with KAPA Pure Beads (Roche, Switzerland), and library amplification was limited to six PCR cycles to reduce amplification bias. After PCR, a final clean-up step was performed to remove any residual primer dimers and non-target fragments. The resulting libraries were assessed for quality and fragment size distribution using a 4150 TapeStation system (Agilent Technologies, USA).

All libraries were then normalized to 20 nM, pooled in equimolar amounts, and subjected to 150 bp paired-end sequencing on an Illumina NovaSeq platform (Illumina, CA, USA). Each library generated approximately 13 million paired-end reads. This deep sequencing strategy provided a comprehensive snapshot of the microbial communities and their genomic characteristics.

#### 2.2.6. Statistics and bioinformatics analysis

The raw fastq sequences were subjected to quality control with TrimGalore^25^. To remove human contaminants, any reads mapping to the human reference genome build 38 with bowtie2^26^ were removed. Taxonomic and functional profiles were calculated via MetaPhlAn 4.0 and HumanN 3.7^27^. Alpha and beta diversities were calculated using the QIIME 2 diversity packages and plotted using custom Python scripts^28^ and the paired t-test was used to identify statistical significance. Differential enrichment analysis was performed using LEfSe^29^. Finally, correlations of microbiome features with metadata were calculated using Spearman correlation with Benjamini-Hochberg correction for multiple testing^30^.

## 3. Results

### 3.1. Cohort overview

In this study, we investigated 17 experienced Polish male mountaineers (23-40 years old) who undertook high-altitude expeditions lasting more than three weeks at altitudes above 3,000 MASL. Before and after their expeditions, we collected comprehensive data including dietary intake, blood and urine parameters, aerobic and anaerobic performance measures, and detailed metagenomic analyses of their gut microbiomes. By integrating these multidimensional datasets, our goal was to characterize how exposure to extreme mountain environments and dietary changes influence gut microbial community structure, functional pathways, and potential health and performance outcomes.

### 3.2. Microbiome changes after the mountaineering expedition

All participants exhibited generally diverse microbiomes both before and after the expedition (Fig. 1a). While the relative abundance of the most plentiful species did not change much in some individuals (A3), others experienced greater shifts in the microbiome (A10). There were on average 200 ± 69 bacterial species identified per participant before and 179 ± 53 species per participant after the expedition. In contrast, we found 76 ± 8 bacterial functions per participant before and 77 ± 8 functions after the expedition. The differences in the numbers of species and functions before and after the expedition were not significant.

**Fig. 1.**
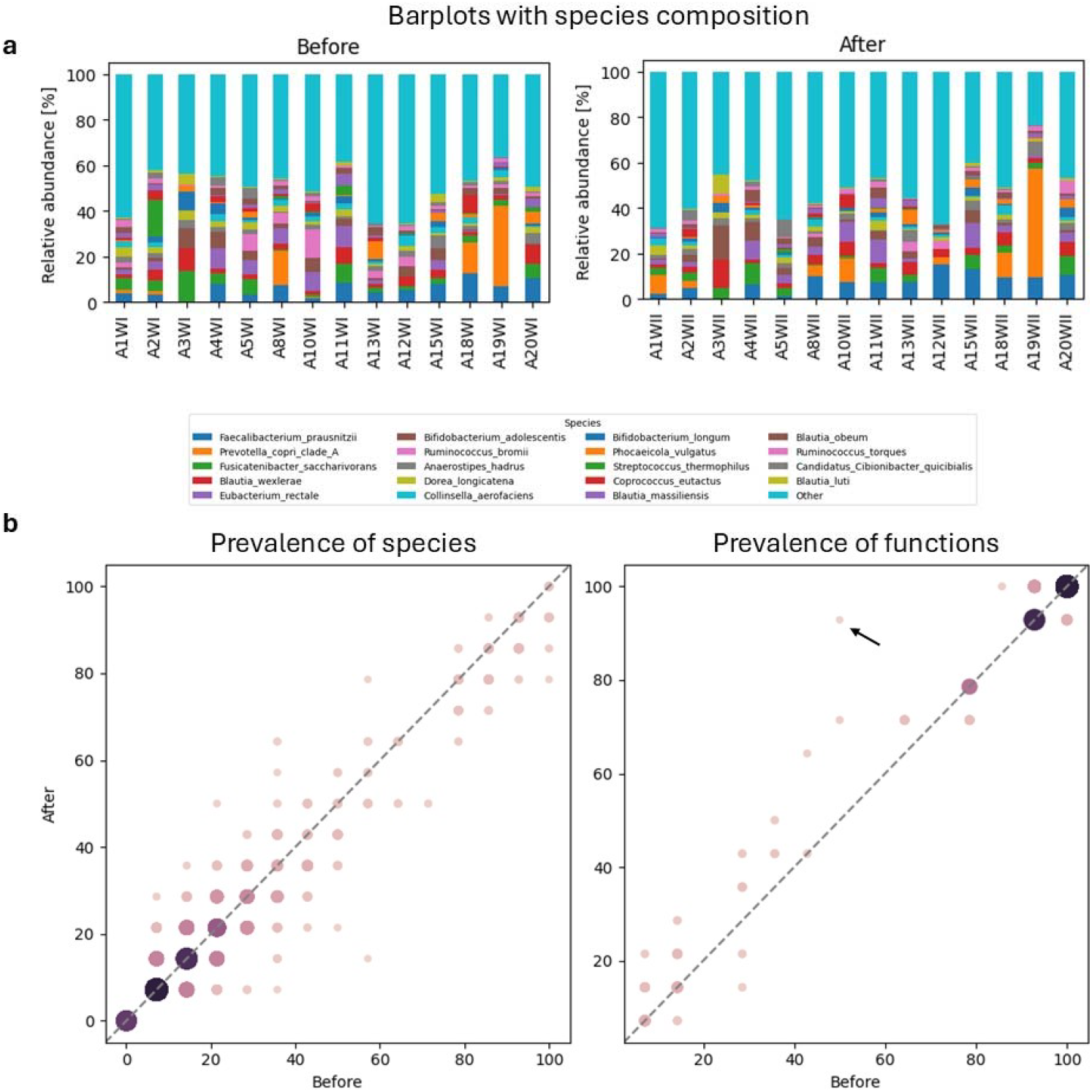
Comparison of gut microbiome compositions before and after the expedition. (a) Barplots with species composition of each sample before and after the expedition. (b) Prevalence of species (left) and functions (right) before versus after the expedition. The *GLUCOSE1PMETAB-PWY*: *glucose and glucose-1-phosphate degradation* pathway, which substantially changed in prevalence as a result of the travel, is marked with a black arrow.

Across all participants, the prevalence of species remained stable after the travel (Fig. ***1***b, left). This was particularly true for species present in less than a third of individuals, and any differences could be attributed to changes in only up to two individuals. We observed a slightly different trend in the comparison of functions - this time, the most stable functions were highly prevalent. There was one outlier, however; it was the *GLUCOSE1PMETAB- PWY: glucose and glucose-1-phosphate degradation* pathway. This function, present in a half of individuals before the expedition, was found in almost everyone upon their return. We observed a statistically significant increase in the abundance of this function as a result of the expedition (Wilcoxon signed-rank test, p<0.05).

A differential enrichment analysis performed using LEfSE revealed two species enriched after the expedition and one before (Fig. 2). The two species which increased in abundance after the expedition were Lactococcus lactis and a species belonging to the Phocaeicola genus. On the other hand, there was a depletion trend of the Clostridiaceae bacterium. We found no statistically significant alterations in microbial functions.

**Fig. 2.**
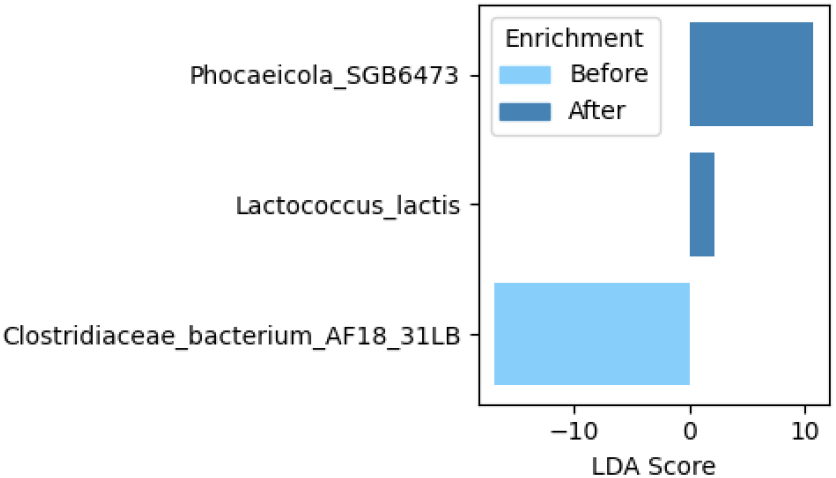
Species enriched in the participants before and after the expedition

Correlating microbiome features with dietary information revealed an association of a *PWY-6703: preQ0 biosynthesis pathway* with vitamin B6 and C intake before the expedition (Spearman correlations of 0.83 and 0.86, respectively, adjusted p-values < 0.05). We observed a decrease in the intake of the vitamins after the expedition, which was significant for vitamin C (vitamin B6: 6.1mg to 2.6mg, T-test p-value > 0.05; vitamin C: 325.7mg to 149.2mg, T-test p–value < 0.01). Similarly, the average abundance of the *PWY-6703: preQ0 biosynthesis* pathway showed a notable, but statistically insignificant decrease (from 18% to 15%). No other associations of functions or species with dietary parameters before or after the expedition passed multiple testing corrections.

### 3.3. Varying degrees of microbiome alterations among participants

Following our observation that the microbiomes were altered in individuals to different degrees (Fig. 1a), we calculated Bray-Curtis distances between matched samples from the same individual, collected before and after the intervention (Fig. 3a). We observed a range of distances between the samples, with samples from the individual A12 being visibly more differentiated than the rest.

**Fig. 3.**
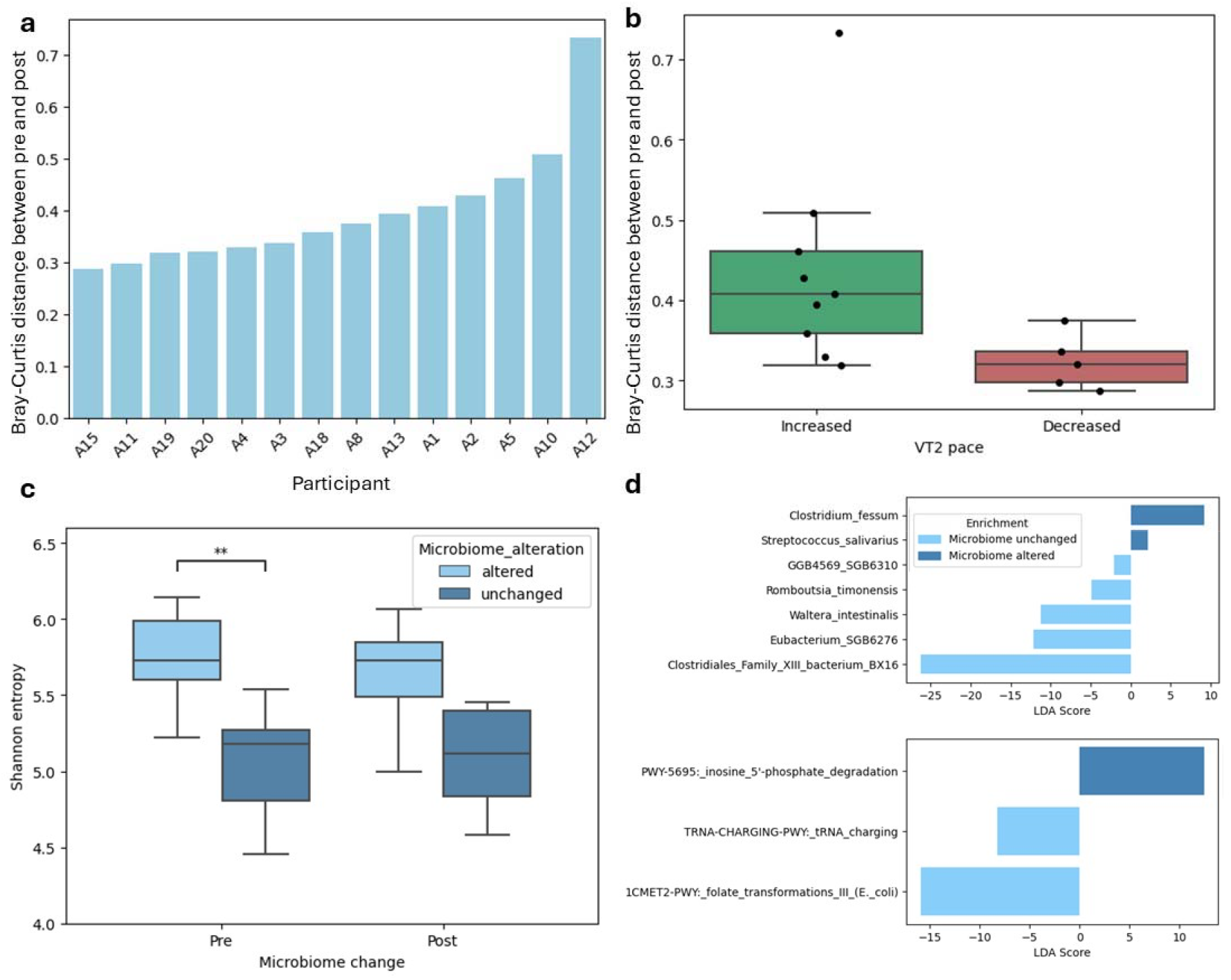
A comparison of participants with greater and smaller gut microbiome alterations after the high-altitude travel. (a) Bray-Curtis distance between matched samples (before and after the expedition) from the same participant. (b) Individuals who observed an improvement in the VT2 pace after the expedition experienced greater microbiome alterations. (c) The microbiome of individuals who experienced greater microbiome alterations was significantly richer before the expedition. (d) LEfSe differential enrichment analysis comparing species and functions in individuals with and without large microbiome alterations at baseline.

Separating the individuals based on their VT2 pace change after the expedition revealed that individuals who experienced an improvement in fitness experienced a notably yet not significantly greater microbiome alteration (Fig. 3b). In addition, participants whose microbiomes changed more (Bray-Curtis distance ≥ 0.36, chosen based on the median) had significantly richer microbiomes before the intervention in comparison to those who experienced subtle shifts (Fig. 3c). This was accompanied by insignificant but notably higher intakes of manganese, calcium, vitamin A, magnesium and phosphorus (mean manganese intake 9.2 and 3.9 mg, calcium 1520.7 and 973.3 mg, vitamin A 2246.0 and 1291 μg, magnesium 687.7 and 397.6 mg, phosphorus 2021.5 and 1545.7 mg in participants with and without substantial gut alterations, p < 0.05, adjusted p-values > 0.05).

A LEfSE differential enrichment analysis of species and functions at baseline revealed statistically significant differences between the two groups (Fig. 3d). The microbiomes of individuals who experienced alterations were enriched in *Clositridium fessum, Streptococcus salivarius* and the *inosine 5’-phosphate* degradation pathway. On the other hand, they appeared to have lower abundances of, among others, a bacterium from the Clostridiales family as well as the *folate transformations* and *tRNA charging* pathways.

### 3.4. Microbiome correlations with blood markers, anaerobic capacity physiological markers and diet in participants exhibiting the strongest microbiome shifts

Investigating the subset of individuals with large microbiome alterations revealed shifts in prevalence of species were more varied than that of functions (Fig. 4a and Fig. 4b). The taxa with the greatest alterations included *Dorea sp AF36_15AT*, present in 57% of individuals before and in 100% after the expedition, as well as *GGB9758_SGB15368* from the Firmicutes phylum, found in 43% before and 86% of individuals after. While the prevalence of functions remained relatively stable, with most functions being present both before and after the expedition, a few functions experienced a substantial prevalence drop. They were *tetrapyrrole biosynthesis I (from glutamate), UDP-N-acetyl-D-glucosamine biosynthesis I* and *O-antigen building blocks biosynthesis (E*.*coli)*, all present in 43% before and 14% of individuals after the expedition.

**Fig. 4.**
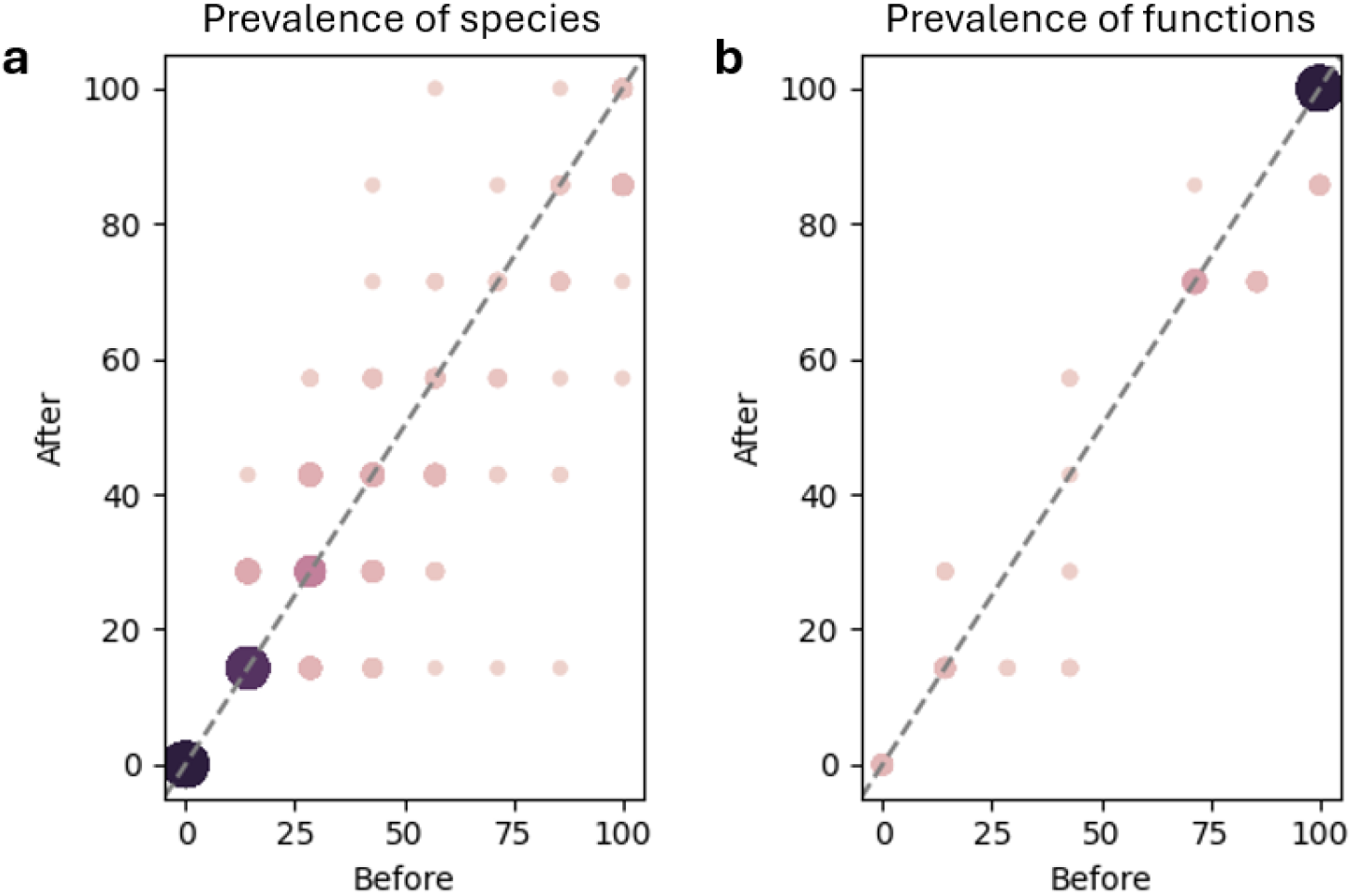
Prevalence comparison of species (a) and functions (b) in individuals with large microbiome alterations as a result of the expedition.

A differential enrichment analysis with LEfSe identified 3 species, namely *Blautia luti, Romboutsia timonensis* and *GGB4569_SGB6310* from the Lactobacillaceae family, enriched in abundance after the expedition. We found no species enriched before, as well as no statistically significant alterations of abundance in functions.

We identified several microbiome feature correlations with blood markers and anaerobic indices (Table 2). Before the expedition, there were two species associated with potassium, specifically *Prevotella copri clade A* (positive correlation) and *Bacteroides uniformis* (negative correlation), as well as 15 cases of marker associations with microbial functions. *dTDP-*β*-L-rhamnose biosynthesis* pathway was the only function associated with two different markers before the expedition, specifically total work performed of lower limbs (W_t_) and mean anaerobic power of lower limbs (P_mean_) (positively in both cases). After the investigation, we found one association with species *Agathobaculum butyriciproducens* with calcium (strong positive correlation) and 8 with functions. This time, two functions were associated with at least two markers (*NAD de novo biosynthesis I (from aspartate)* positively with basophils and calcium; and *superpathway of L-serine and glycine biosynthesis I* negatively with sodium and positively with cholesterol HDL). Finally, two functions had different associations both before and after the expedition. G*uanosine ribonucleotides de novo biosynthesis* was positively correlated with sodium before and negatively with mean platelet volume (MPV) after, while *myo-, chiro- and scyllo-inositol* degradation was positively associated with sodium before and with basophils after.

**Table 2.**
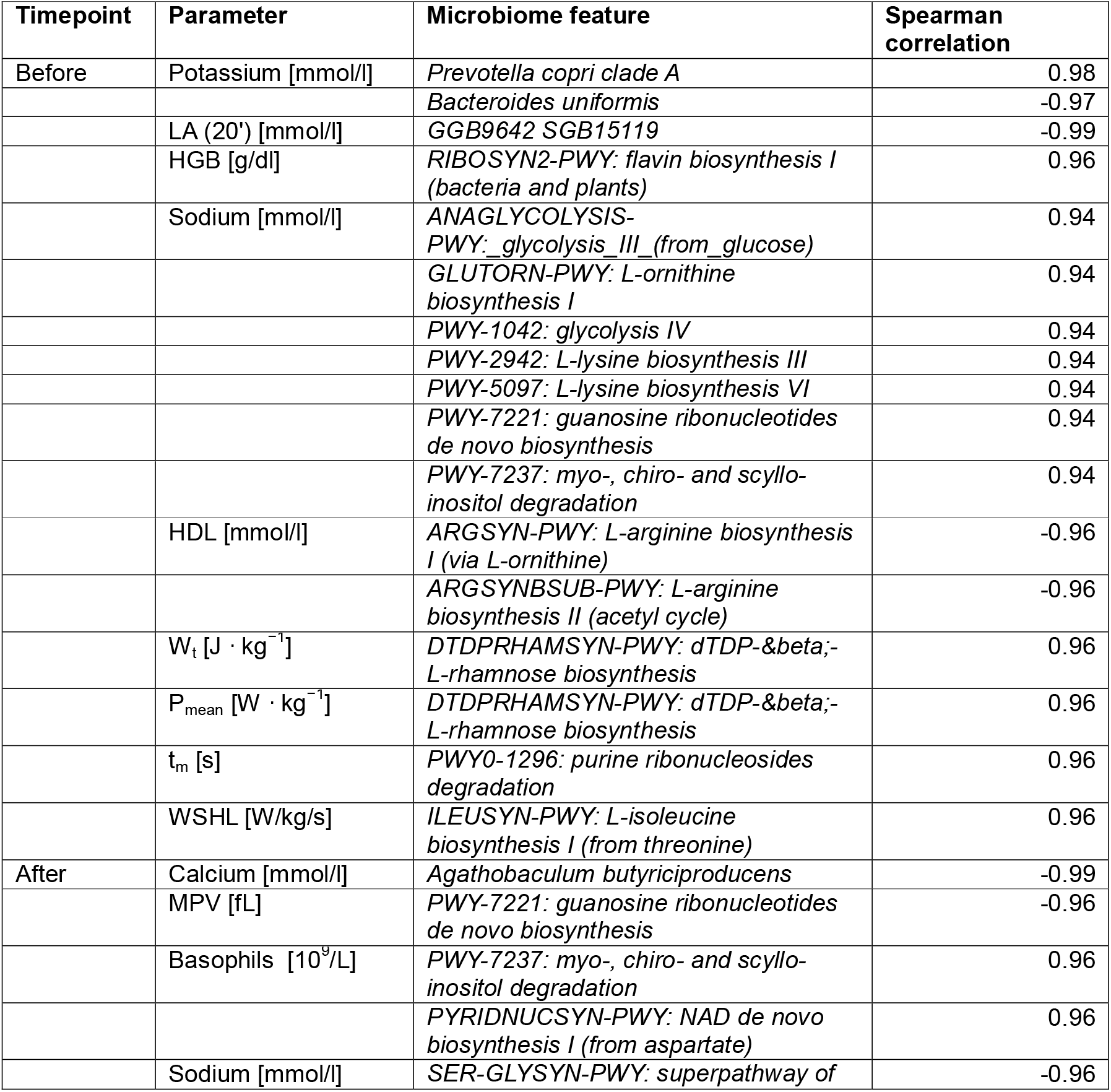
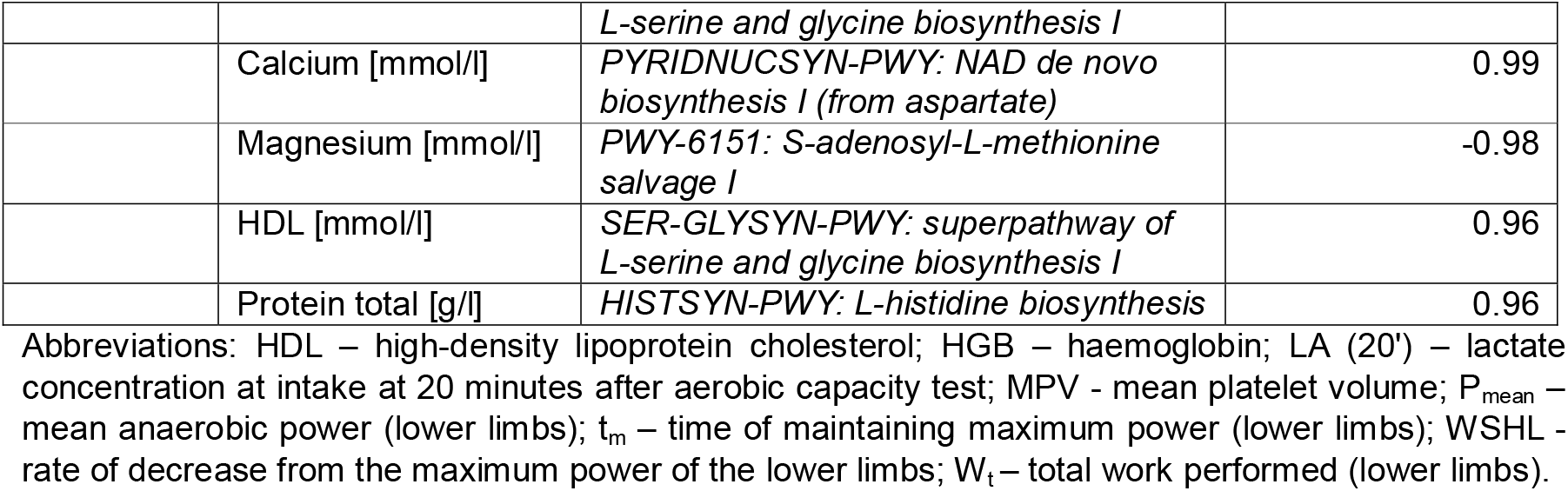
Statistically significant correlations (adjusted p-values < 0.05) of blood markers and anaerobic indices with microbiome features in individuals with microbiome alterations, before and after the expedition (n=7).

Finally, we found a number of microbial function associations with diet after the expedition (all of which passed the correction for multiple testing, Supplementary Table 1). We identified strong positive correlations (Spearman correlation > 0.90) of *coenzyme A biosynthesis* with overall calorie and carbohydrate intake, monounsaturated fatty acid (MUFA) as well as vitamins C, B12, B1 and B6. *L-arginine biosynthesis* was associated with overall carbohydrate intake and digestible carbohydrates, while the *penthose phosphate* pathway with selenium and plant protein with *sucrose biosynthesis*. Negative correlations were found between *adenine and adenosine salvage* and sodium, as well as between *L-rhamnose biosynthesis* and animal protein.

## 4. Discussion

### 4.1. Microbiome and Nutritional Adaptations to High-Altitude Stress

The observed changes in gut microbiome compositions before and after the high-altitude mountaineering expedition indicate adaptive shifts in the gut microbiome and host physiology in response to extreme environmental and physiological stressors. Our findings demonstrate that exposure to low atmospheric pressure, hypoxic conditions, and challenging dietary changes can reshape the gut microbiome, potentially affecting host metabolism, immune function, and overall performance. Similar to previous studies showing that altitude-induced hypoxia alters energy metabolism and physiological adaptations in athletes^31–33^, we identified functional shifts in the gut microbiome, such as increased prevalence of the *glucose and glucose-1-phosphate degradation* pathway, which was present in half of individuals before the expedition and in almost all participants upon their return. In addition, the abundance of this pathway increased significantly. The observed increase in the prevalence of the *glucose and glucose-1-phosphate degradation* pathway and the variable enrichment of specific bacterial taxa indicate a microbiome fine-tuning to optimize energy utilization under adverse conditions.

From a nutritional perspective, our results highlight the importance of vitamin intake, particularly vitamins C and B6, in modulating microbial metabolic pathways such as the *preQ0 biosynthesis* pathway. The observed association between the *preQ0 biosynthesis* pathway and dietary intake of vitamins B6 and C prior to the high-altitude mountaineering expedition highlights the intricate interplay between nutrition, microbiome dynamics, and metabolic adaptation. Vitamin B6, known for its role as a cofactor in amino acid metabolism^34^, is essential for the synthesis of nucleotides and neurotransmitters. This vitamin may enhance the efficiency of metabolic processes involved in preQ0 production, as the pathway is likely reliant on amino acid precursors. Additionally, vitamin C serves as a potent antioxidant, crucial for mitigating oxidative stress that can be exacerbated in high-altitude environments^35–37^. Its antioxidant properties may also support beneficial gut microbial populations capable of metabolizing this vitamin and subsequently influencing pathways like the preQ0 biosynthesis pathway. The dietary intake of these vitamins may modulate gut microbiome composition, ensuring the availability of substrates necessary for microbial metabolism, which in turn supports the *preQ0 biosynthesis* pathway. This association underscores the significance of adequate vitamin intake in preparing the body for the metabolic demands of high-altitude exposure, where increased energy expenditure and altered oxygen availability necessitate robust cellular adaptations^38^.

An analysis of Bray-Curtis distances between baseline and post-expedition samples revealed varying degrees of microbiome alterations among the participants. Individuals with greater microbiome shifts responded to the high-altitude conditions better through improvements in fitness markers such as a higher pace at the VT2. They also appeared to have significantly richer baseline microbiomes in comparison to individuals with subtle shifts. This was accompanied by insignificant but notably higher intakes of manganese, calcium, vitamin A, magnesium and phosphorus before the expedition. Finally, their microbiomes were enriched in *Clositridium fessum, Streptococcus salivarius* and the inosine 5’-phosphate degradation pathway.

Before the expedition, species-level associations such as *Prevotella copri* and *Bacteroides uniformis* with potassium, along with pathway associations like *dTDP-*β*-L-rhamnose biosynthesis* linked to physical performance markers, suggested a baseline connection between microbial composition, electrolyte balance, and energy metabolism in participants with large microbiome alterations. Post-expedition shifts in functional pathways indicated gut microbiome associations with immune markers and lipid metabolism. Dynamic connections, such as the reversal of *guanosine ribonucleotide biosynthesis* and *inositol degradation* pathway correlations, imply the microbiome’s role in modulating nutrient metabolism, immune response, and electrolyte regulation.

A differential enrichment analysis with LEfSe in our study identified 3 species, namely *Blautia luti, Romboutsia timonensis* and *GGB4569_SGB6310*, enriched in abundance after the expedition. Similarly, Su et al. showed a relative and absolute increase in *Blautia A* abundance in men who were exposed to high altitudes (>3,600 MASL) for more than 2 months^39^. *Blautia A* species were shown to be involved in a module of “cobalamin biosynthesis” and butyric acid production that are both beneficial to microbial ecosystems and intestinal epithelial cells^39,40^. An *in vivo* animal experiment supported a crucial role of *Blautia A* species in facilitating host fitness to hypoxia environments, likely via anti-inflammation and intestinal barrier protection to maintain intestinal health, thereby suggesting a high translational potential of *Blautia A* species as a candidate probiotic agent for the prevention or treatment of hypoxia-associated maladaptation or disorders. Taken together, these observations of increased abundance of *Blautia A* imply this genus is a beneficial high-altitude bacterial group that may play an important role in promoting acclimatization and adaptation to high-altitude conditions^39^.

We observed significant microbial functional associations with dietary intake after the expedition. Notably, strong positive correlations with *coenzyme A biosynthesis* and various dietary components suggest that these nutrients may enhance microbial metabolic pathways essential for energy production and lipid metabolism. Similarly, positive associations of *L-arginine biosynthesis* with overall carbohydrate intake and the assimilation of carbohydrates indicate that dietary carbohydrates may support microbial synthesis of this amino acid, which is crucial for various physiological functions^19,41,42^. Furthermore, associations of the *pentose phosphate* pathway with selenium and plant protein with sucrose biosynthesis highlight the potential role of specific micronutrients and macronutrients in modulating microbial metabolic pathways. On the other hand, negative associations of *adenine and adenosine salvage* pathways with sodium intake, as well as between *L-rhamnose biosynthesis* and animal protein consumption could suggest competitive inhibition or substrate limitation in these metabolic processes. Overall, the results emphasize complex interactions between dietary components and microbial metabolic activities in the gut. However, the sample size for this analysis was limited (n=7); thus, future validation of these findings in a larger cohort is necessary.

### 4.2. Practical Implications, Limitations, and Future Perspectives

From an applied perspective, these findings provide valuable guidance for mountaineers, coaches, and sports nutritionists. Pre-expedition strategies might include optimizing micronutrient status, particularly for vitamins and minerals shown here to correlate with beneficial microbial functions and performance markers. During expeditions, integrating nutrient-dense, microbiome-supportive foods – even if partially in the form of supplements or freeze-dried products – may help maintain beneficial gut microbiota profiles and associated metabolic functions. Beyond the athletic context, our findings contribute to a growing body of literature on the role of gut microbiome in adaptation to extreme environments^39,43–48^, highlighting opportunities to leverage nutritional interventions to support both health and performance.

Future research should aim to confirm these findings in larger cohorts and explore targeted interventions-such as prebiotics, probiotics, postbiotics, or tailored micronutrient supplementation to promote a beneficial gut microbiome response in high-altitude environments. Integrating metagenomics with host transcriptomics, metabolomics, and immune profiling, as well as conducting controlled feeding studies, could further elucidate the mechanisms linking the gut microbiome, diet, and physiological adaptation in extreme sports and beyond.

Our study has some limitations. The climbers did not constitute a single group, led by a single climbing goal in the territory of a single country and mountain range. Climbers, characterized by the characteristics described in the study, constitute a small but elite group in Poland, hence several distinct expedition groups, climbing at similar altitudes and with similar sporting goals, but in different parts of the world, qualified for the study. Gathering and encouraging participants was one of the most challenging stages of the project.

In addition to the level of hypoxia, other factors such as nutrition and the type of physical activity performed in the mountains, as well as the number of active days versus the number of rest days, may contribute significantly to the observed changes in microbiome. Climbing groups active in Peru could have used the downtime between several days of mountain activity to descend to a town at the foot of the mountains to compensate for the negative energy balance there, while expedition groups in Kyrgyzstan, Nepal or Pakistan were dependent on the food they took into the mountains or that the expedition agency offered them.

## 5. Conclusions

In conclusion, our study demonstrates that the gut microbiome is responsive and potentially malleable to the dietary and environmental stresses encountered during high-altitude mountaineering. The observed changes in microbial composition and function, their associations with micronutrient intake, and their potential links to improved athletic performance underscore the integral role of the microbiome as an adaptive partner in extreme human endeavors^49^.

## Data availability

The raw fastq sequences have been deposited to the European Nucleotide Archive (ENA) under the accession PRJEB84244.

## Ethics approval

This study was performed in line with the principles of the Declaration of Helsinki. Approval was granted by the Bioethics Committee of the Cracow Regional Medical Chamber (11/04/2022, No. 68/KBL/OIL/2022).

## Authors’ contributions

EKG, BF and KHL conceptualized the study; EKG, KZ and KHL conducted the data collection and processing; EKG, KHL and KZ produced metadata; KZ, PPŁ and TK performed the statistics and bioinformatics analysis; EKG and BF obtained funding; EKG and BF provided of study materials; EKG and KHL developed methodology; KZ visualized the data; EKG, BF and KHL managed the project; BF and KHL supervised the project; EKG, KZ and KHL drafted the manuscript. All authors edited the manuscript and contributed to its final version. All authors have read and approved the final version of the manuscript, and agree with the order of presentation of the authors.

