## Supplementary Table 1 for "High-altitude mountaineering induces adaptive gut microbiome shifts associated with dietary intake and performance markers": Supplementary Table 1.docx

Running heading: Adaptive gut microbiome shifts in high-altitude mountaineers

Ewa Karpęcka-Gałka^1,*^, Kinga Zielińska^2^, Barbara Frączek^3^, Paweł Łabaj^2^, Tomasz Kościółek^4^, Kinga Humińska-Lisowska^5^

^1^ Doctoral School of Physical Culture Sciences, University of Physical Culture in Krakow, 31-571 Krakow, Poland

^2^ Malopolska Centre of Biotechnology, Jagiellonian University, 30-387 Krakow, Poland

^3^ Department of Sports Medicine and Human Nutrition, Institute of Biomedical Sciences, University of Physical Culture in Krakow, 31-571 Krakow, Poland

^4^ Sano Centre for Computational Medicine, 30-054 Krakow, Poland

^5^ Faculty of Physical Education, Gdansk University of Physical Education and Sport, 80-336 Gdansk, Poland

*Corresponding author:

**Supplementary information**

**Supplementary Table 1.** Statistically significant correlations (adjusted p-values < 0.05) of energy and nutritional value of the daily food ration with microbiome features in individuals with microbiome alterations, after the expedition.

| **Parameter** | **Microbiome feature** | **Spearman correlation** |
| --- | --- | --- |
| Energy [kcal] | *COA-PWY-1:_superpathway_of_coenzyme_A_biosynthesis_III* | 0.96 |
| Vitamin C [mg] | *COA-PWY-1:_superpathway_of_coenzyme_A_biosynthesis_III* | 0.96 |
| Vitamin B12 [µg] | *PWY-7851:_coenzyme_A_biosynthesis_II* | 0.96 |
| Vitamin B6 [mg] | *PWY-7851:_coenzyme_A_biosynthesis_II* | 0.96 |
| Vitamin B1[mg] | *PWY-7851:_coenzyme_A_biosynthesis_II* | 0.96 |
| Digestible carbohydrates [g/kg_mc] | *ARGSYN-PWY:_L-arginine_biosynthesis_I_(via_L-ornithine)* | 0.96 |
|  | *ARGSYNBSUB-PWY:_L-arginine_biosynthesis_II_(acetyl_cycle)* | 0.96 |
| Selenium [µg] | *NONOXIPENT-PWY:_pentose_phosphate_pathway_(non-oxidative_branch)_I* | 0.96 |
| Sodium [mg] | *PWY-6609:_adenine_and_adenosine_salvage_III* | -0.96 |
| MUFA [g] | *COA-PWY:_coenzyme_A_biosynthesis_I_(prokaryotic)* | 0.96 |
| Animal protein [g] | *DTDPRHAMSYN-PWY:_dTDP-&beta;-L-rhamnose_biosynthesis* | -0.96 |
| Vegetable protein [g] | *PWY-7238:_sucrose_biosynthesis_II* | 0.96 |

Abbreviations: MUFA - monounsaturated fatty acids
